# Dissecting sequence determinants of DNA methylation and in silico perturbation

**DOI:** 10.1101/2025.11.20.689274

**Authors:** Faming Chen, Qinhao Zhang, Shiwei Zhu, Jinxia Liang, Siyun Zhong, Zehong Zhang, Jianzhong Su, Yangfan Guo, Xiaoying Fan, Yan Zhang, Fulong Yu

## Abstract

DNA methylation shapes cellular identity and genome function, yet the sequence logic that governs its establishment remains largely unresolved. Here we present MethylAI, a cross-species–pretrained and human-specialized deep learning framework that predicts single-CpG methylation states directly from genomic sequence with high fidelity. Through quantitative attribution, MethylAI uncovers intrinsic cis-regulatory principles embedded in DNA, revealing conserved transcription factor (TF) motifs whose influence extends beyond CpG composition. Activated motifs demarcate gene-body CpG islands, encode lineage-defining methylation states, and align with TF occupancy and diverse histone modifications, thereby linking sequence syntax to local chromatin regulation. CTCF perturbation experiments validated MethylAI-predicted methylation shifts at activated motifs, showing strong concordance with in silico perturbations. Mapping genetic variants onto methylation-linked active motifs further uncovered widespread connections to zinc-finger TF families, providing mechanistic insight into how noncoding variation contributes to human traits and diseases. By transforming DNA sequence into interpretable methylation grammar, MethylAI establishes a generalizable framework for decoding the regulatory architecture of the human epigenome.

## Introduction

The genetic code provides the blueprint of life, yet its interpretation depends on epigenetic modifications that shape how DNA is read and utilized^1^. DNA methylation, the most pervasive and conserved of these modifications, plays a defining role in transcriptional regulation, development, and cellular identity^2,3^. It establishes stable yet reversible regulatory states across the genome, and its perturbation is a hallmark of developmental disorders, cancer, and aging^4,5^. Despite decades of investigation, the rules by which genomic sequence dictates DNA methylation remain only partially understood. Early studies identified CpG density and GC content as strong correlates of methylation levels^6^, giving rise to the classical concept of CpG islands, which are frequently hypomethylated and enriched for regulatory activity^7,8^. Yet these features reflect only a narrow aspect of a far more intricate regulatory logic.

In practice, DNA methylation is shaped by a coordinated molecular machinery in which DNA methyltransferases and demethylases act together with transcription factors, histone modifiers, and chromatin remodelers to establish and maintain context-specific methylation states^7,9^. These processes operate as an integrated system that senses, interprets, and reinforces sequence-encoded regulatory cues, linking local DNA composition to chromatin accessibility, transcriptional output, and higher-order genome organization. Across cell types and species, this interplay produces dynamic yet reproducible methylation landscapes^10^, implying the existence of a deeply encoded sequence grammar governing methylation regulation. Learning this grammar in a generalizable manner and enabling its systematic dissection in silico represents a critical step toward understanding the principles that shape the human epigenome.

To address this challenge, we developed MethylAI, a deep learning framework that learns how genomic sequence encodes methylation states across evolution and cellular contexts. Through large-scale cross-species pretraining followed by human-specific fine-tuning, MethylAI accurately predicts single-CpG methylation directly from DNA sequence while preserving interpretability at nucleotide resolution. This framework enables systematic identification of sequence determinants that govern methylation, reveals previously unrecognized transcription factor signatures that bridge sequence composition with epigenetic state, and illuminates how genetic variants perturb methylation by disrupting these encoded rules. To promote transparency and community use, we provide the complete codebase, pretrained models, and harmonized methylation compendium as publicly available resources.

## Results

### Comprehensive sequence-based DNA methylation prediction using MethylAI

DNA methylation follows dynamic evolutionary trajectories across species and exhibits strong cell type–specific and context-dependent variation^3,11^. We hypothesized that accurate prediction of methylation directly from sequence, coupled with quantitative attribution of individual nucleotides, could reveal intrinsic cis-regulatory determinants of methylation variation. By systematically dissecting sequence features that shape methylation signals, we aimed to deconvolve the underlying sequence-encoded logic.

To capture both conserved and divergent principles of methylation regulation, we assembled the largest cross-species methylation compendium to date, comprising 1,900 single-CpG–resolved whole-genome bisulfite methylomes across 12 species (Extended Data Fig. 1a, Supplementary Table 1). Leveraging this resource, we developed MethylAI, a framework built upon large-scale cross-species pretraining followed by human-domain fine-tuning, enabling high-accuracy prediction of human methylation patterns solely from genomic sequence (Fig. 1a, Extended Data Fig. 2). During pre-training, we extracted 18-kb CpG-centered sequences from reference genomes, totaling 25.17 billion CpGs and 463.93 trillion nucleotides. Species-specific batch normalization and adaptive output heads (Extended Data Fig. 3) enabled stable learning across phylogenetically distant species and heterogeneous tissue sources. Building on this foundation, we designed multi-scale convolutional modules that preserve intermediate features at distinct receptive fields (Fig. 1a, Extended Data Fig. 3), allowing the model to integrate both local motifs and distal sequence dependencies. The resulting contextual embeddings spontaneously organized into clusters corresponding to functional genomic categories without explicit supervision (Fig. 1b), indicating that the model learns biologically meaningful sequence representations. We further incorporated a multi-task objective to jointly predict single-site methylation and regional methylation averages across 200 bp, 500 bp, and 1 kb windows, reflecting spatial coherence of methylation along the genome.

**Fig. 1.**
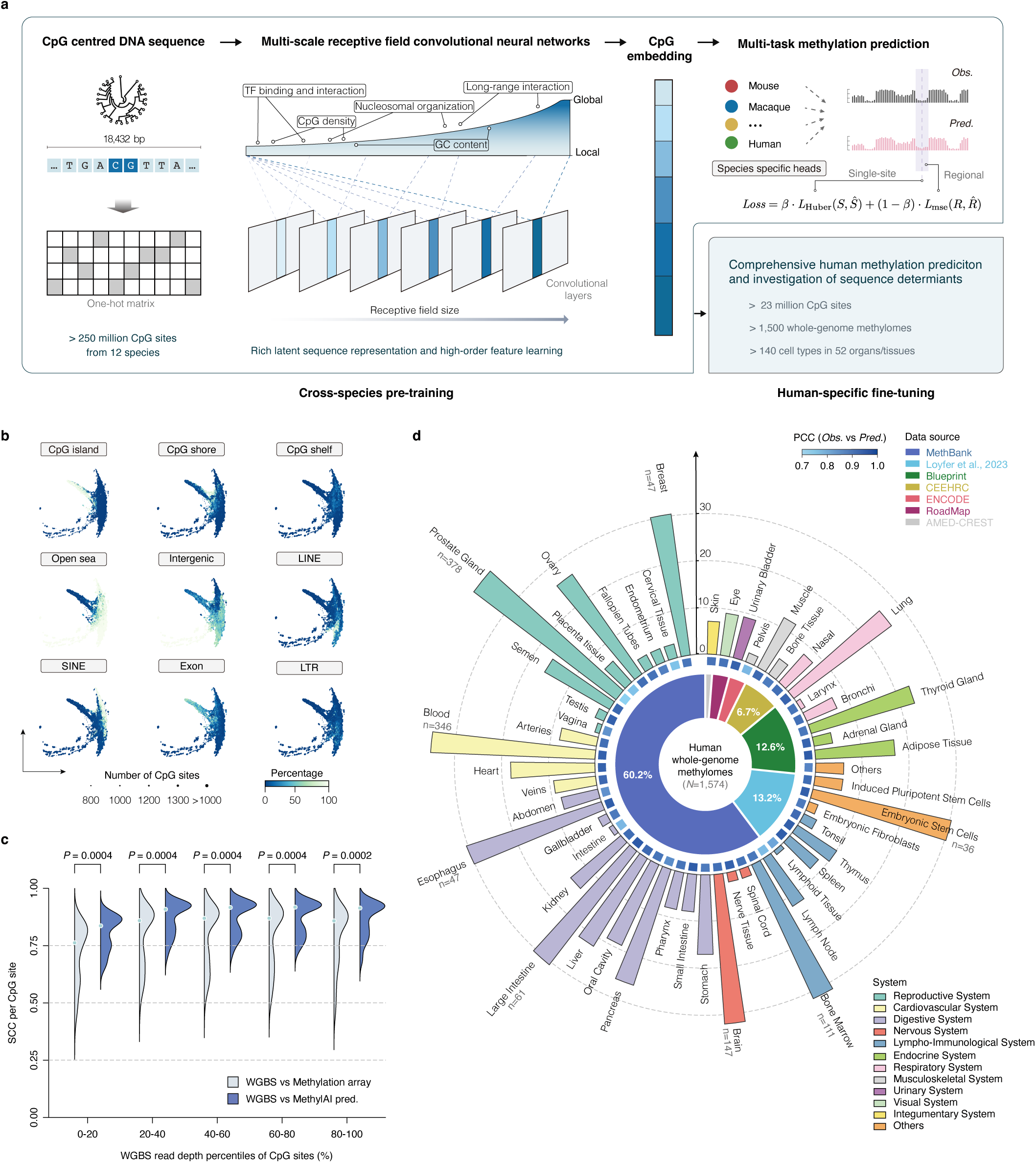
Overview of the MethylAI architecture and predictive performance. a, Schematic of the model architecture, illustrating cross-species pre-training and human-specific fine-tuning of MethylAI. Massive CpG-centered sequences from 12 species were encoded and taken as input into multi-scale receptive field convolutional neural networks, which capture effective features at different levels, both locally and globally, such as transcription factor (TF) binding sites, CpG density, nucleosome organization, and GC content. Representations of CpG-centered sequences are learned at different scales and integrated; these are designated as ‘CpG embeddings’. Methylation prediction tasks for both single-sites and regions were designed as training objectives. b, Multi-scale potential of heat-diffusion for affinity-based trajectory embedding (PHATE) projection of CpG embeddings, presented in separate panels for distinct genomic annotations (CpG island, shore, shelf, open sea, intergenic, LINE, SINE, exon, LTR). Each point represents a coarse-grained cluster of CpG sites, with point sizes indicating the number of CpGs represented and color denoting the percentage of methylation within each element. This illustrates the relative distributions of genomic functional elements within the embedding space. c, Violin plots depicting the Spearman correlation coefficient (SCC) between whole-genome bisulfite sequencing (WGBS) data and MethylAI predictions (light blue), as well as between WGBS and methylation array data (light grey), stratified by WGBS read depth percentiles at CpG sites (x-axis). Each violin represents the distribution of SCCs within a depth bin, with overlaid dots indicating the mean. P-values were calculated using two-sided Wilcoxon signed-rank tests. d, Catalog of human whole-genome methylomes included in this study (n = 1,574). The inner pie chart shows proportions by major organ/system; the middle heatmap displays the Pearson correlation coefficient (PCC) between observed (highest depth sample per tissue/type) WGBS methylation and MethylAI predictions; the outer ring presents sample counts per category, with reference ticks at N = 10, 20, 30. Bars exceeding 30 are compressed and labeled with their exact counts. Data sources are identified in the panel (MethBank, Loyfer et al., 2023; Blueprint; CEEHRC; ENCODE; RoadMap; AMED CREST); a complete list of samples and statistics is provided in the Methods and Supplementary Tables 1.

Human-specific fine-tuning utilized 1,574 whole-genome methylomes spanning 238 cell types across 52 tissues and 12 physiological systems. Using one high-coverage representative sample per tissue, MethylAI achieved a mean Pearson correlation coefficient (PCC) of 0.903 (range: 0.741–0.962) on an independent test dataset (Fig. 1d). Ablation analyses demonstrated that cross-species pretraining was the dominant performance determinant, substantially outperforming alternatives such as doubling the input sequence length (Extended Data Fig. 4).

We next benchmarked MethylAI against INTERACT^12^, a CNN–Transformer hybrid for methylation prediction, and HyenaDNA^13^, a genomic foundation model (Methods). MethylAI outperformed both across the genome and in diverse genomic contexts, achieving a mean PCC of 0.875 compared with 0.813 for INTERACT and 0.828 for HyenaDNA across 1,574 test samples under matched model sizes (Extended Data Fig. 3). This advantage extended to comparison with experimental methylation arrays using WGBS as ground truth: MethylAI achieved mean correlation scores of 0.854–0.923 across coverage categories, exceeding the 0.803–0.871 range observed for arrays (Fig. 1d, Supplementary Table 2). These results thus establish MethylAI as a robust and accurate framework for sequence-based DNA methylation prediction, achieving high fidelity across diverse tissues and cell types using only genomic sequence information.

### The sequence composition and patterns contributing to the methylation

Equipped with MethylAI’s high-fidelity predictions, we next examined the sequence determinants underlying DNA methylation. We computed base-resolution attribution scores using DeepSHAP^14^, which estimate the marginal contribution of each nucleotide to the model’s prediction (Methods), thereby quantifying both activating and repressive influences on methylation levels in lung tissue. For representative analysis, we randomly sampled 100,000 CpG sites, given the computational demands of profiling all ~27 million CpGs in the human genome. For each CpG, MethylAI quantified the contribution of every nucleotide within the input sequence. Attribution decayed sharply with distance from the focal site: 31.7% of total contribution arose from the nearest ±1 kb region, a 5.8-fold enrichment relative to the mean contribution across the full sequence (Fig. 2a). We therefore designated this ±1-kb window as the primary analytical scope for subsequent analyses.

**Figure 2.**
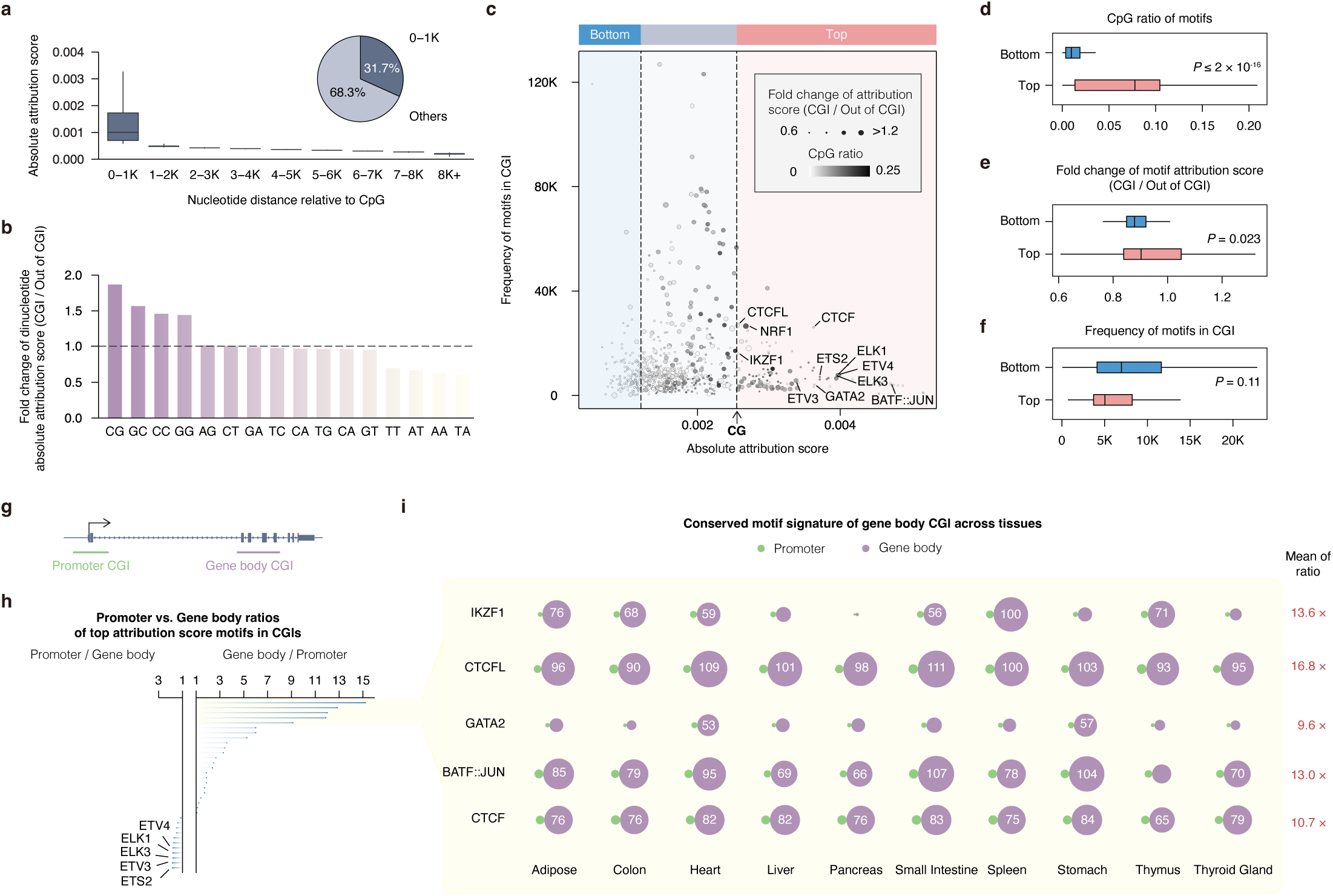
Tissue-conserved TF signature distinguishes gene-body from promoter CGIs. a, Sequence contribution analyzed by distance from CpG sites. The boxplot (left) shows model-derived absolute attribution scores binned by nucleotide distance (kb). The pie chart (right) displays the proportion of total attribution scores originating within 1 kb (31.7%) versus beyond (68.3%). b, Bar plot showing the fold change of dinucleotide absolute attribution scores inside versus outside CpG islands (CGI), calculated as the ratio of mean attribution scores (CGI/non-CGI) and ordered by magnitude. The dashed line at 1.0 indicates no change. c, Scatterplot comparing absolute mean motif attribution scores (x-axis) with motif frequency (y-axis, in thousands) within CGIs. Each point represents a motif, where point size encodes the attribution score fold change (CGI vs. non-CGI) and color intensity reflects the CpG ratio (darker = higher). The dashed line marks the contribution score of the CG dinucleotide itself. Motifs are classified as “Top” (light red) or “Bottom” (light blue) based on their position relative to this line. d–f, Statistical comparisons between “Top” and “Bottom” motif groups within CGIs. The “Top” group exhibits a significantly higher CpG ratio (d, P ≤ 2 × 10^−16^), a greater fold-change in attribution scores (e, P = 0.023), and a non-significant difference in motif frequency (f, P = 0.11). g, Schematic distinguishing promoter CGIs (green) from gene-body CGIs (purple) relative to gene architecture. h, Horizontal bar plot showing the relative enrichment of active motifs in promoter versus gene-body CGIs. Ratios are calculated to highlight enrichment in either region (e.g., promoter/gene-body or vice versa). The top five promoter-enriched motifs are labeled. i, Venn diagram detailing the tissue-level distribution of the top five gene-body-enriched motifs. Circle area is proportional to the number of motif instances (counts ≥ 50 are shown). The right column reports the mean gene-body/promoter ratio per tissue. For all boxplots (a, d–f), the center line indicates the median, the box represents the interquartile range (IQR; 25th–75th percentiles), and whiskers extend to 1.5×IQR from the box edges; outliers are not shown. P-values in d–f were derived from two-sided Wilcoxon signed-rank tests. Detailed calculations are available in the Methods.

We sought to assessed how single nucleotides and dinucleotides contribute to methylation prediction. Cytosine (C) and guanine (G) consistently showed elevated attribution values, comparable to dinucleotides containing at least one C or G, whereas the CpG dinucleotide displayed a markedly stronger contribution (P = 2.2 × 10^−16^; Extended Data Fig. 6). This quantitatively recapitulates the well-established role of CpG composition in shaping methylation patterns^6^.

We next asked whether larger sequence motifs further influence local methylation states. Although numerous TFs have been implicated in methylation regulation^9,15–17^, the extent and specificity of their genome-wide effects remain unclear. Treating TF binding motifs as coherent regulatory units allows us to interrogate their contribution systematically. From 781 TF motif profiles precomputed across the human genome, we selected significant motif instances (P < 1×10^−4^) for evaluation. Strikingly, 19.5% of these motifs were more informative for methylation prediction than the CpG dinucleotide itself (Extended Data Fig. 6), suggesting that TF binding features exert broader and more substantial influence over methylation patterns than previously appreciated. Notably, motif presence alone did not predict attribution magnitude, indicating that local sequence context and combinatorial interactions shape motif activity. The most frequently occurring motifs belonged to zinc-finger (ZNF) families (e.g., ZNF384, ZNF135, ZNF281, ZNF213, ZNF460, ZNF148, Extended Data Fig. 6). Ranking motifs by median absolute attribution scores identified strong contributors to methylation regulation, including YY2, AP-1 complex members (FOS, FOSB, FOSL2, JUN, JUNB, JUND), CTCF, GATA2, and ELK4, underscoring their substantial regulatory roles (Extended Data Fig. 6).

### MethylAI reveals tissue-conserved TF signatures of gene body CGIs

Motivated by these findings, we sought to investigate TF signatures underlying CpG islands (CGIs), the most prominent regulatory elements associated with DNA methylation. Conceptually, CGIs are defined by sequence features of high GC content and elevated CpG frequency^7,18^ and typically exhibit hypomethylation across cell types^7,8^. Owing to their well-established sequence characteristics, CGIs provide an ideal context for validating known sequence determinants and uncovering new features captured by the model. We analyzed attribution scores for CGIs shorter than 1 kb and compared them with randomly selected CpG sites described above. Among all dinucleotide combinations, the CG dinucleotide consistently showed the highest attribution score within CGIs and a 1.87-fold higher score than in non-CGI regions (Fig. 2b, Extended Data Fig. 7a). This pattern quantitatively recapitulates the central role of CpG density in defining CGI features and regulatory functions.

Most CGIs remain constitutively hypomethylated in somatic cells, however, the potential contribution of TF binding motifs to CGI methylation patterns has not been systematically investigated. Using MethylAI, we identified 128 TF motifs (16.4% of all examined) within CGIs whose mean absolute attribution scores exceeded that of the CG dinucleotide, including well-established methylation-associated factors such as CTCF and NRF1^16,19^ (Fig. 2c, Extended Data Fig. 7a). Ranking motifs by average attribution score revealed that the highest-scoring motifs exhibited markedly greater CpG ratio in their sequence composition than the lowest-scoring ones (P < 2×10^−16^; Fig. 2d). In contrast, the two groups showed no significant difference in occurrence frequency (P = 0.11; Fig. 2f) and only a modest difference in attribution fold change between CGI and non-CGI regions (P = 0.023; Fig. 2e). These results indicate that a high CpG ratio is the principal distinguishing feature of top motifs within CGIs, likely reflecting the intrinsically CpG-rich sequence context of these elements. Notably, exceptions such as BATF::JUN and GATA2 maintained strong attribution despite low CpG ratios, whereas ELK1, ETV3, and ETV4 combined high CpG density with high attribution (Fig. 2c), suggesting both CpG-dependent and CpG-independent contributions to CGI regulation.

To assess whether TF motifs contribute differentially across CGI subclasses, we partitioned CGIs into promoter-associated and gene-body–associated categories (Fig. 2g). ETS family motifs (e.g., ETS2, 1.86-fold enrichment) were preferentially enriched in promoter CGIs, consistent with their established role in promoter-proximal regulation^20,21^. In contrast, IKZF1 (15.20-fold), BATF::JUN (11.88-fold), CTCF (9.13-fold), and related motifs showed strong enrichment in gene-body CGIs (Fig. 2h). Remarkably, this motif architecture was highly conserved across ten additional tissues: the top five motifs (IKZF1, CTCFL, GATA2, BATF::JUN, CTCF) consistently exhibited 9.6–16.8-fold enrichment in gene-body CGIs (Fig. 2i). Genes containing CTCF-anchored active motifs within their gene-body CGIs were functionally enriched for calcium ion binding, cell–cell adhesion, and plasma membrane components (Extended Data Fig. 7d), a pattern reproduced across tissues (Extended Data Fig. 7c). Collectively, these findings reveal a conserved and previously underappreciated role of the gene body–anchored TF motifs in shaping CGI methylation landscapes and suggest potential mechanisms for maintaining chromatin organization across diverse tissues.

### Motif-specific activation underlying cellular identities

Uniquely unmethylated regions (UURs) are methylation-defined elements that mark cell identity–specific loci, exhibiting restricted hypomethylation in contrast to the pervasive hypomethylation of CGIs. They are enriched for enhancer-associated binding sites of lineage-determining TFs^9,22,23^. We compiled previously defined UUR sets across 39 human cell types^24^ and selected the top 1,000 per cell type for downstream analysis. A dedicated MethylAI model fine-tuned on this dataset, with targeted oversampling of CpGs in UURs (Methods), achieved high predictive fidelity at these loci (average PCC = 0.924; Extended Data Fig. 8).

To quantify motif activation as a property reflecting both the strength of sequence attribution and the fidelity of motif-TF correspondence, we defined a motif activation score (MAS). MAS integrates the average attribution score with motif-match likelihood and is normalized by motif length, capturing both the sequence’s impact on methylation and its similarity to established TF binding profiles. This formulation supports direct comparison across motifs with heterogeneous lengths and PWM structures. The MAS reliably distinguished functional motifs: although KLF17 matched its known PWM, it was not activated, whereas ETV1 and RUNX1 showed strong, T-cell-specific activation (Fig. 3a).

**Fig. 3.**
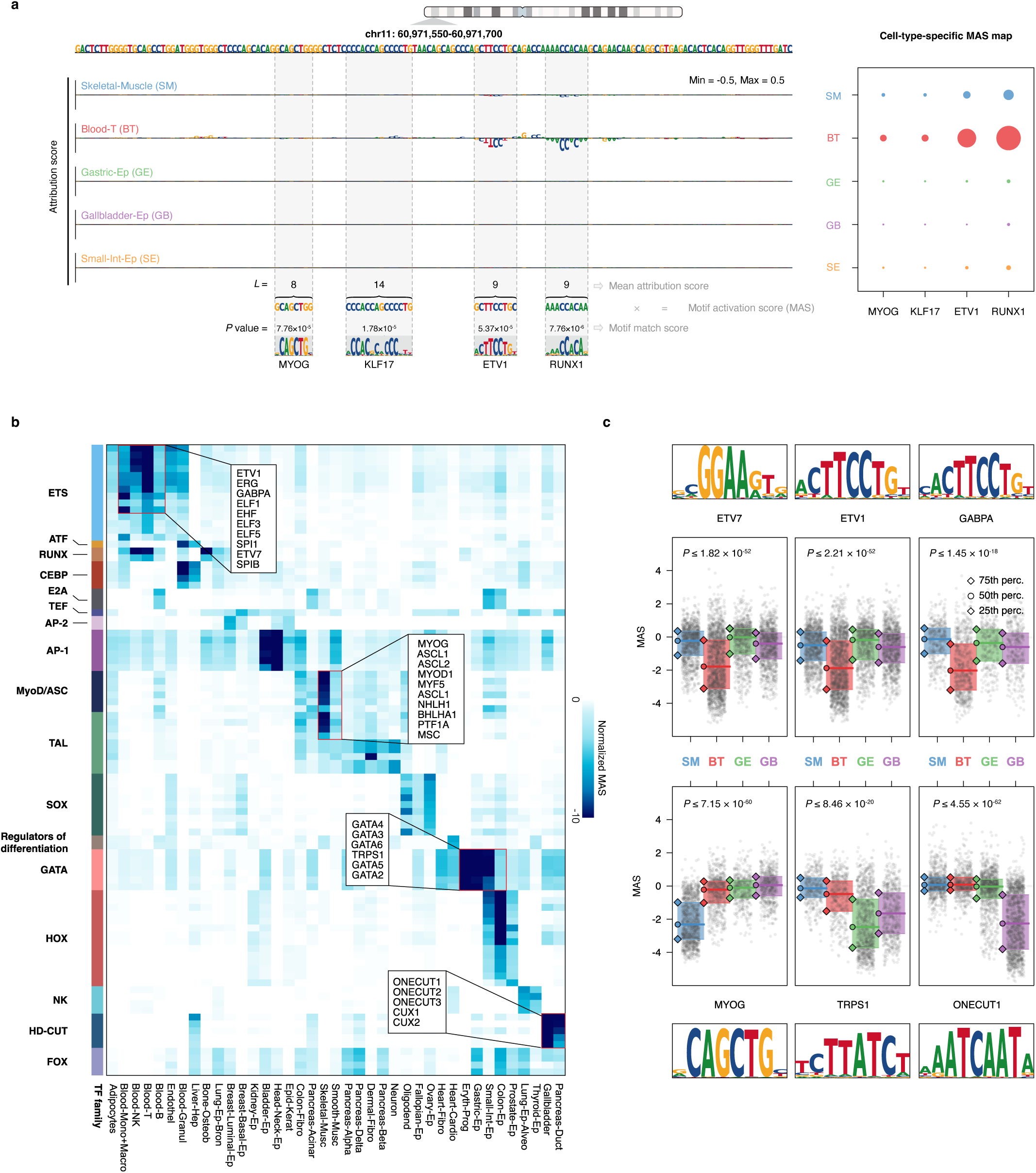
TF motif activation specifies cellular identities a, Example locus (chr11:60,971,550-60,971,700) illustrating the calculation of motif activation. Left: Per-base attribution tracks (MethylAI gradient saliency) for selected tissues. For each nucleotide, letter height reflects its attribution score (scale: −0.5 to 0.5). Shaded boxes highlight motif instances with their length (L) and PWM match score. Right: Bubble plot of the tissue-level motif activation score (MAS) for selected motifs. Bubble area is proportional to the MAS, and color indicates the tissue (SM, Skeletal Muscle; BT, Blood T; GE, Gastric Ep; GB, Gallbladder Ep; SE, Small Int Ep). The per-instance ActivationScore is defined as the PWM match score multiplied by the mean attribution across the motif window; these scores are then averaged to yield the final MAS. b, Heatmap of normalized MAS values for TF families across tissues, computed on a curated set of Uniquely Unmethylated Regions (UURs, n = 1,246). Rows (motifs/families) and columns (tissues) are hierarchically clustered to reveal tissue-specific activation patterns. Color intensity reflects normalized activation strength (values ≤ −10 are clipped for display). Major motif families are annotated in the side color bar. c, Boxplots showing the distribution of MAS for selected motifs across four tissues, with individual regional scores shown as jittered grey points. Motif logos are displayed above each panel. Plotted values are the signed log1p-transformed MAS (sign(x)·ln(1+|x|)). For each motif, statistical comparisons were performed between the two tissues with the lowest aggregate MAS. Attribution scores are derived from MethylAI gradient saliencies. Further details on all calculations, normalization, UUR definition, and clustering parameters are provided in the Methods. For boxplots, the center line is the median, the box represents the interquartile range (IQR), and whiskers extend to 1.5×IQR. P-values were computed using two-sided independent-samples t-tests on untransformed MAS, and values below display precision are noted with ‘≤’.

In addition, we systematically compared MAS values for all TF motifs across UURs from the 39 human cell types. The resulting motif activation maps revealed sharply defined cell-type specificity, which became even more pronounced after normalization (Methods), aligning closely with established biological knowledge (Fig. 3b). Master regulators exhibited clear lineage-dependent activation patterns: ETS family motifs showed stronger negative MAS in immune cells (monocytes, NK cells, T cells, and B cells)[PMID:26301869], whereas MyoD/ASC and TAL family motifs displayed stronger negative signals in skeletal muscle cells^25^ (Fig. 3b). Representative TF motifs such as ETV7, ETV1, GABPA, MYOG, TRPS1, and ONECUT1 showed the expected lineage-specific MAS profiles (Fig. 3c). Hence, these motif activation patterns define cell-type-specific regulatory signatures that delineate lineage programs encoded in the methylation landscape.

### Motif activation corresponds to functional chromatin engagement

Although DNA methylation is catalyzed by DNA methyltransferases (DNMTs) and ten-eleven translocation (TET) enzymes, its establishment and maintenance occur within the architecture of chromatin. Chromatin accessibility, histone modification landscapes, and TF occupancy collectively shape where methylation machinery can engage DNA, thereby defining local regulatory states^7^. To determine how motif activation learned by MethylAI relates to these broader chromatin features, we integrated experimental epigenomic profiles and protein interaction networks.

We first performed WGBS on HEK-293T cells and identified 91,705 hypomethylated regions (Methods). Among these, 52,538 (57.3%) contained at least one active motif as defined by MAS. Prominent motif families included GATA factors (e.g., GATA6), AP-1 complex components (e.g., JUN), and zinc-finger-related families (e.g., BNC2, CTCF) (Fig. 4a). Notably, the motif for REST, a transcriptional repressor known to interact with methylation machinery^16^, displayed the highest average MAS across all families (Fig. 4a).

**Fig. 4.**
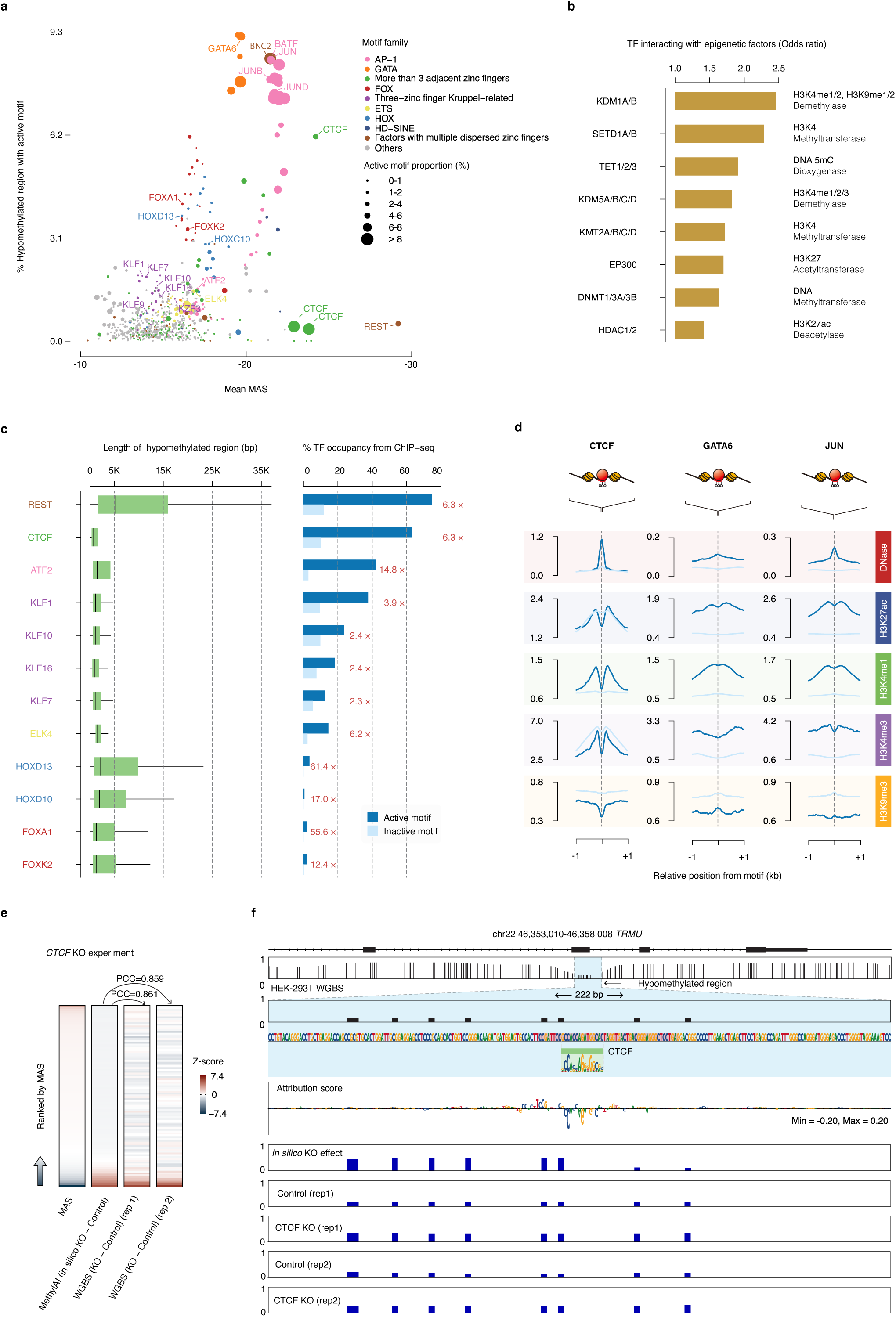
Motif activation reflects chromatin and methylation regulation validated by perturbation experiments. a, Scatterplot correlating the mean motif activation score (MAS) with the proportion of hypomethylated regions regulated by each motif in HEK-293T cells (n = 91,705 motifs). Point area is proportional to the percentage of regulated regions, and color indicates the motif family. b, Enrichment analysis (odds ratio) of physical interactions between TFs of the most active motifs (top 30% by regulated regions) and major epigenetic protein families. Epigenetic regulator categories are listed on the right. c, Comparison of active (dark blue) and inactive (light blue) motifs. Left: Boxplots of hypomethylated region length. Right: Bar plot showing TF binding, measured as the percentage of motifs overlapping with corresponding ChIP-seq peaks (ENCODE data). Fold enrichment is annotated. d, Metaplots showing average signal profiles of DNase-seq and active histone marks (H3K27ac, H3K4me1, H3K4me3) centered on active (dark blue) and inactive (light blue) motifs for CTCF, GATA6, and JUN. Lines represent mean signal intensity. Note that y-axis scales differ between plots. e, Heatmap comparing predicted methylation changes from in silico CTCF knockout with observed changes from experimental WGBS data. Values are Z-score normalized. The Pearson correlation between prediction and experiment is shown. f, Genome browser view of the TRMU locus. Tracks include gene annotation, experimental WGBS data (control and CTCF KO), an active CTCF motif, and the corresponding MethylAI attribution and in silico KO prediction tracks. In c, boxplots show the median (center line), interquartile range (IQR; box), and 1.5×IQR (whiskers); outliers are not shown. Further details on motif definitions and all calculations are provided in Methods.

To assess whether active motifs are biochemically linked to methylation and chromatin pathways, we examined protein–protein interactions between the corresponding TFs and epigenetic regulators. TFs associated with the top 30% of motifs (ranked by their frequency in hypomethylated regions) showed substantially more interactions with DNA methylation and chromatin regulators (odds ratio = 1.42–2.46) than other TFs (Fig. 4b). These interactions spanned DNMT and TET enzymes, H3K4 methyltransferases and demethylases, and histone acetylases and deacetylases (Fig. 4b), indicating that these TF families broadly interface with chromatin remodeling and methylation pathways.

We next evaluated whether these active motifs are associated with TF binding experimentally. Using HEK-293/HEK-293T TF and histone ChIP-seq data from the ENCODE project, we found that active motifs were more frequently overlapped by TF ChIP-seq peaks than inactive motifs (Fig. 4c), indicating a strong correspondence between predicted motif activation and experimentally observed TF occupancy.

Chromatin features surrounding these motifs further supported their functional relevance. Motifs for CTCF, ATF2, KLF1/7/10/16, and ELK4 preferentially resided within shorter hypomethylated regions, whereas motifs for REST, HOXD13, and HOXC10 occurred within longer regions (Fig. 4c). Active motifs for GATA6 and JUN were associated with elevated chromatin accessibility (DNase-seq signal; Fig. 4d) and strong active histone marks (H3K27ac, H3K4me1, H3K4me3; Fig. 4d), accompanied by reduced repressive H3K9me3 signals (Fig. 4d). Specifically, active CTCF motifs displayed stronger DNase and H3K4me1 signals, reduced H3K4me3 and H3K9me3 signals, and comparable H3K27ac levels relative to their inactive counterparts (Fig. 4d), consistent with the distinct chromatin-anchoring role of CTCF^26^. These active motifs do not merely correlate with hypomethylation but are embedded within open, transcriptionally engaged chromatin marked by TF binding and activating histone modifications. Thus, although trained solely on methylation data, MethylAI implicitly learns higher-order chromatin features and regulatory interactions that shape methylation dynamics.

### Experimental validation of transcription factor regulation in DNA hypomethylation

To experimentally validate the regulatory role of TFs predicted by MethylAI in shaping DNA hypomethylation, we performed targeted knockout experiments in HEK-293T cells, followed by whole-genome DNA methylation profiling. We first focused on CTCF, a zinc-finger protein known to bind chromatin insulators and maintain hypomethylated domains.

Using the MethylAI in silico perturbation framework, we predicted methylation changes upon CTCF depletion. These predictions showed strong concordance with experimentally observed methylation shifts after CTCF knockout (PCC = 0.859–0.861; Fig. 4f), confirming that MethylAI accurately identifies regions whose methylation state depends on CTCF binding.

To further test causality at the motif level, we deleted a predicted CTCF-active motif located within an exon of the *TRMU* gene using CRISPR/Cas9. Loss of this motif led to a marked increase in local DNA methylation (Fig. 4e), demonstrating that the hypomethylated state at this locus is directly maintained by CTCF occupancy. Together, protein-level and motif-level perturbations validate that CTCF binding is essential for sustaining hypomethylation at a subset of regulatory regions identified by MethylAI.

### MethylAI predicts genetic effects on DNA methylation

Understanding how genetic variants influence DNA methylation is fundamental to linking genotype with regulatory outcomes. Genetic perturbations to methylation can alter chromatin architecture, gene expression, and ultimately phenotype, yet comprehensive characterization remains limited by cohort size and experimental throughput. Classical methylation quantitative trait locus (mQTL) studies have identified millions of variant–CpG associations, with up to 45% of CpG sites under cis genetic control^27,28^. However, these analyses interrogate only measured loci and cannot explore the extensive sequence space of unobserved variants. A model capable of predicting methylation changes induced by sequence variation would allow in silico reconstruction of mQTL landscapes and provide mechanistic insight into how genetic variation modulates methylation regulation across tissues.

To assess this capability, we benchmarked MethylAI’s in silico variant predictions against the multi-tissue mQTL atlas from the Enhancer GTEx (eGTEx) project^29^, the first comprehensive mQTL map across solid tissues. We extracted all significant cis-mQTL CpG sites (mCpGs; FDR < 0.05) and their associated variants (P < 1×10^−5^), yielding 6,525,489 mQTL–mCpG pairs across seven tissues. Predicted methylation changes from MethylAI showed strong correspondence with observed allelic effects, with larger predicted perturbations associated with greater absolute slope values (Extended Data Fig. 10a). MethylAI correctly predicted the direction of methylation change for more than 95% of the 20,000 most significant GTEx mQTLs, and maintained high accuracy across tens of thousands of additional variants (Extended Data Fig. 10b). Prediction accuracy was highest when mQTLs were derived from the same tissue as the model input, underscoring the tissue specificity of methylation regulation (Extended Data Fig. 10b).

We next examined mQTLs located within tissue-specific active motifs identified by MethylAI, incorporating both statistically significant and conditionally independent variants. In these regions, MethylAI achieved 87.0–89.8% accuracy in predicting the direction of methylation change—substantially higher than for mQTLs in inactive (53.2–58.1%) or non-motif (52.6–57.9%) regions across ±1 kb windows (Fig. 5a). Predicted effect magnitudes and experimentally derived absolute slopes were also significantly greater for mQTLs residing in active motif regions (Extended Data Fig. 10c), indicating stronger functional coupling between sequence perturbation and methylation outcome at these loci.

**Fig. 5.**
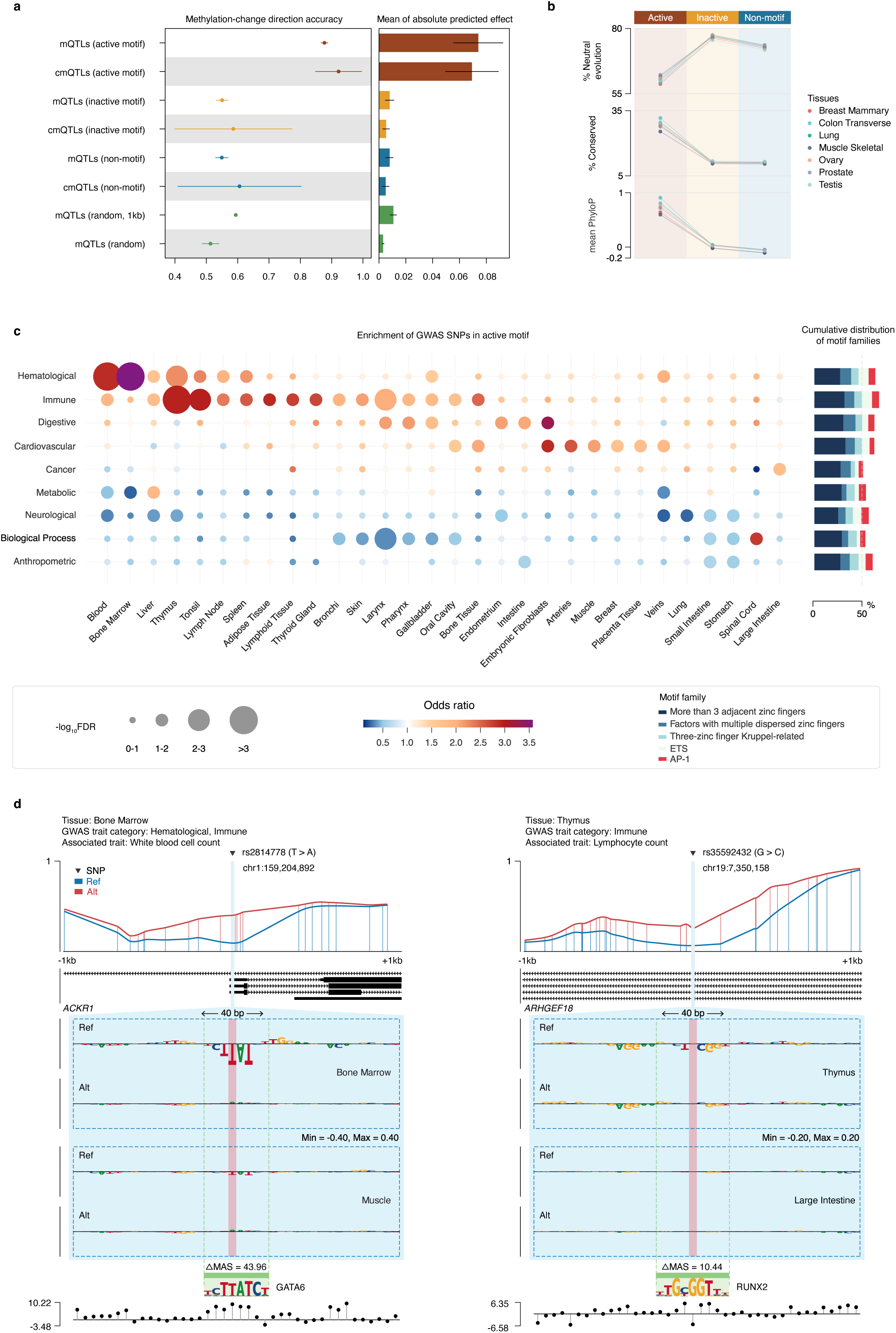
MethylAI predicts genetic effects on DNA methylation in silico. a, Prediction accuracy for mQTL effect direction (left) and predicted effect size (right) for all mQTLs and conditionally independent mQTLs (cmQTLs). Performance is evaluated across different genomic contexts: active motifs, inactive motifs, non-motif regions, and random genomic windows. Points show mean values, and error bars represent standard deviation (SD) across 7 tissues. b, Evolutionary metrics for different region classes. Plots show (from top to bottom): percentage of regions under neutral evolution, percentage under conservation, and mean 470-way PhyloP score. Regions are classified as neutral (−1 ≤ PhyloP ≤ 1) or conserved (PhyloP > 1). Colors correspond to tissues as labeled. c, GWAS trait enrichment in tissue-specific active motifs. Left: Bubble plot showing enrichment odds ratio (color) and statistical significance (−log₁₀(FDR); area). Non-significant associations (FDR > 0.1) are shown as minimal-sized bubbles. Right: Stacked bar plot showing the distribution of top associations across major TF motif families for each trait. d, Genome browser views of two example GWAS loci: rs2814778 (ACKR1, associated with white blood cell count) and rs35592432 (ARHGEF18, associated with lymphocyte count). Tracks shown (top to bottom) include: allele-specific MethylAI predictions, GENCODE gene annotations, MethylAI attribution scores, motif locations, and allele-specific ΔMAS.

To assess evolutionary constraint at these regulatory hotspots, we evaluated phyloP conservation across genomes from 470 mammalian species^30^. Tissue-specific active motif regions exhibited significantly elevated conservation relative to other genomic regions (Fig. 5b). These conserved, motif-anchored loci therefore represent hotspots where evolutionary constraint converges with functional genetic influence on DNA methylation, marking them as key nodes in hereditary methylation control.

### Mapping motif activation underlying human traits and diseases

To evaluate whether tissue-specific active motifs contribute to human complex traits and disease risk, we systematically assessed the correspondence between GWAS variants and motif activation states. Across genome-wide hypomethylated loci from 52 human tissues, 36,768 GWAS-associated SNPs (approximately 10% of all variants in the GWAS Catalog) overlapped either active (1,686 SNPs) or inactive (35,082 SNPs) motif regions. These variants represented 1,232 distinct traits spanning eight major phenotypic categories. For each trait category, we quantified tissue-specific enrichment of SNPs within active motifs using inactive motifs as background.

Clear tissue-relevant enrichment patterns emerged (Fig. 5c). Hematopoietic traits were enriched in blood and bone marrow motifs; immune traits in tonsil, thymus, adipose, lymphatic, and thyroid tissues; cardiovascular traits in arterial and fibroblast lineages; and liver-associated traits in hematopoietic, immune, and metabolic tissues. Beyond these tissue-specific enrichments, trait-associated variants showed a striking convergence on shared TF motif families: nearly half of GWAS variants located in active motifs mapped to zinc-finger (ZNF) binding sites, including motifs with adjacent, dispersed, or three-zinc-finger Kruppel domains (Fig. 5c). This pattern is consistent with the broad involvement of ZNF proteins in tissue homeostasis and diverse human diseases^31^, suggesting that many disease-associated variants converge on conserved ZNF-centered regulatory mechanisms despite their tissue-specific manifestation.

To dissect the molecular consequences of risk variants in active motifs, we used MethylAI to predict the direction and magnitude of methylation changes within ±1 kb of 1,039 variants with defined risk alleles (Supplementary Table 5). Across these loci, 83.4% of risk alleles were predicted to increase local methylation (866 of 1,039), accompanied by widespread attenuation of motif activation (828 of 866). Representative examples highlight tissue-anchored, motif-centered mechanisms.

For hematopoietic traits, the rs2814778 risk allele in the 5′UTR of ACKR1, a locus linked to white blood cell count, was predicted to substantially increase methylation at nearby CpGs (maximum predicted effect = 0.21; Fig. 5d). This SNP disrupts a GATA6 motif embedded within an evolutionarily conserved bone marrow region (REF MAS = −46.24 vs. ALT MAS = −2.28), consistent with the essential roles of GATA factors in erythroid and megakaryocytic regulatory programs^32^.

For immune traits (basophil, eosinophil, and lymphocyte counts), the risk allele of rs62111672 mapped to a RUNX2 motif within the ARHGEF18 gene body, active in thymus (REF MAS = −12.01 vs. ALT MAS = −1.57). The risk allele was predicted to increase methylation (maximum effect = 0.34; Fig. 5e), consistent with RUNX2’s nuclear expression in infiltrating immune cell subsets and its role in immune response initiation^33^.

For metabolic traits such as type 2 diabetes, the risk allele of rs72729610 in the TRIM2 gene body was predicted to elevate local methylation by disrupting an ONECUT3 motif active in liver (REF MAS = −56.41 vs. ALT MAS = −14.52; maximum effect = 0.52). ONECUT3 is known to regulate metabolic transcriptional programs including glycolysis^34^. By systematically modeling variant-induced methylation perturbations across tissues and motif contexts, MethylAI reveals a regulatory landscape in which trait-associated variants act through tissue-specific, motif-centered mechanisms that link sequence variation to human phenotypes and diseases.

## Discussion

We developed MethylAI, a sequence-based deep learning framework that predicts DNA methylation at base-pair resolution with high accuracy and interpretability. Through large-scale cross-species supervised pretraining followed by human-domain specialization, MethylAI captures evolutionarily conserved sequence rules that generalize across diverse genomic contexts. The model integrates multi-scale feature extraction and complementary prediction tasks at both single-CpG and regional levels, enabling it to represent local sequence syntax and long-range regulatory dependencies simultaneously. This design allows MethylAI not only to recapitulate known principles of methylation regulation but also to uncover previously unrecognized sequence determinants and transcription factor signatures that shape methylation variation across tissues and species.

A key advance of MethylAI lies in transforming our understanding of how DNA sequence encodes methylation potential quantitatively. Previous studies described sequence dependence of methylation mainly through qualitative features such as GC content or CpG density, which offered only limited mechanistic insight. In contrast, MethylAI adopts a data-driven framework that learns sequence and methylation relationships directly from billions of CpG-centered instances across species without relying on prior assumptions about sequence composition. This enables a fully quantitative, base-level measurement of how each nucleotide contributes to methylation outcomes within its genomic context. The model refines canonical features like GC richness into precise and continuous effects and further uncovers the quantitative influence of less explored elements including transcription factor motifs and higher order sequence combinations. Such data-driven interpretability provides a generalizable foundation for evaluating any hypothetical or newly discovered sequence feature across diverse contexts, turning DNA methylation from an observational pattern into a computable regulatory function.

Importantly, sequence-to-function models such as MethylAI can act as biological oracles, translating the static language of DNA into functional readouts that guide experimental design. This capability opens new opportunities for rational epigenetic engineering. For instance, the in silico design of synthetic enhancers or promoters with desired methylation states or cell-type-specific activities. Such predictive control could enable next-generation synthetic biology, where regulatory elements are engineered to maintain stable epigenetic configurations or dynamically respond to environmental cues.

Although MethylAI is ultimately designed for interpreting and discovering human methylation sequence dependence, its large-scale pretrained model and learned embeddings provide a versatile foundation for a wide spectrum of methylation-related studies. The pre-trained representations can serve as an effective initialization or feature backbone for downstream applications, enabling rapid adaptation to new tasks with limited data. While genome-scale language models such as HyenaDNA have advanced sequence understanding, our results demonstrate that task-specific pre-training on methylation signals confers a clear advantage for methylation-related inference. Lightweight fine-tuning and transfer learning based on MethylAI unlock unprecedented potential across multiple domains, including developmental methylation dynamics, early disease detection, aging prediction, single cell methylome analysis, and translational research.

In sum, MethylAI transforms DNA sequence into a quantitatively interpretable map of methylation logic and provides a scalable foundation for mechanism modeling and new biology discovery. As experimental throughput and modeling capacity continue to grow, such sequence-to-function frameworks will bridge prediction with control, guiding the next generation of causal, programmable, and design-oriented epigenomics.

## Methods

### Assembly of multi-species whole-methylomes

#### Data collection

We collected whole-genome bisulfite sequencing (WGBS) data from public databases including ENCODE, IHEC, and MethBank (Supplementary Table 1). The compiled dataset comprises 1,900 samples across 12 species, of which 1,574 are human samples spanning 52 tissues and 238 cell types (Supplementary Table 1). For each sample, we extracted the sequencing coverage and the number of reads supporting methylation for each CpG site from processed files. Data from forward and reverse strands were merged when available.

#### Quality control

We applied the following quality control (QC) pipeline. For ENCODE and IHEC data, we first retained only autosomal CpG sites, excluding those on sex chromosomes (X and Y) and sites within 18,432 bp of chromosome ends (to avoid edge effects in model input). CpG sites with sequencing coverage below five were classified as low-quality. Samples with over 50% of CpGs failing this threshold were discarded. For human samples from MethBank, which aggregates data from diverse studies and exhibits greater quality heterogeneity, we applied two additional quality controls using the provided metrics: samples with bisulfite conversion rates below 97% or genomic coverage below 10× were excluded.

#### Preprocess

We computed raw site-level methylation values and smoothed methylation levels using the bsmooth function from R package bsseq^35^ (with parameters: ns = 35, h = 500). Regional methylation levels were derived by averaging raw methylation values across consecutive, genomic windows of 1 kb, 500 bp, and 200 bp.

For training, validation, and test sets, we excluded any CpG site that was low-quality in more than 50% of samples. This filtering was applied only to the modeling datasets, while all CpGs were retained for downstream analyses.

For non-human species, all autosomal CpGs were used in training. For human data, autosomes were partitioned as follows: chromosomes 1-9 and 12-22 were used for training; chromosome 10 was used for validation; and chromosome 11 was reserved for testing. This scheme was consistently applied during both pretraining and fine-tuning.

### MethylAI pre-training and fine-tuning

#### MethylAI architecture

MethylAI was implemented in PyTorch. The input is a one-hot encoded DNA sequence of length 18,432 centered on a target CpG site (encoding: A = [1,0,0,0], T = [0,1,0,0], G = [0,0,1,0], C = [0,0,0,1]). The model outputs an S × 5 vector representing five methylation metrics for *S* samples: raw site methylation, smoothed site methylation, and regional methylation levels for three window sizes (200 bp, 500 bp, and 1 kb).

The model consists of three main components: an input block, six multi-scale convolution blocks, and a species-specific output block. The input block includes a convolutional layer followed by species-specific batch normalization and an exponential linear unit (ELU), then two repeated stacks of convolution, species-specific batch normalization, and GELU activation. Each multi-scale convolution block contains two convolutional sub-blocks, each composed of two residual-connected stacks of convolution, species-specific batch normalization, and GELU. Key hyperparameters of multi-scale convolution blocks include input/output channels, depth (number of repeats), kernel size, and stride. The species-specific output block consists of a crop layer, adaptive average pooling, a linear layer, ELU activation, a final linear layer, and a sigmoid activation.

#### MethylAI pre-training and fine-tuning

MethylAI was first pretrained on the multi-species WGBS dataset and then fine-tuned on human dataset. During pretraining, all autosomal CpG sites from non-human species were used. For human data, chromosomes 1–9 and 12–22 were used for training, chromosome 10 for validation, and chromosome 11 for testing.

During pretraining, each batch was alternately sampled from one of the 12 species. Training proceeded for 140,000 steps (each step = one batch from each species), totaling 1,680,000 batches over two epochs. We used the AdamW optimizer with a learning rate of 6×10^−4^ and a batch size of 300 (50 per GPU across 6 GPUs). A warmup schedule was applied for the first 30% of the first epoch, linearly increasing the learning rate from 0 to 6×10^−4^, followed by a constant learning rate. In the second epoch, a cosine annealing scheduler reduced the rate from 6×10^−4^ to 6×10^−7^. Loss functions were task-specific: mean squared error (MSE) for smoothed site and regional methylation, and Huber loss for raw site methylation. Specifically, to mitigate the influence of low-confidence data, we ignored the loss from low-quality CpG sites. This was implemented by reading the sequencing coverage for each CpG site in every sample and setting the loss to zero for sites with coverage less than five. Finally, all losses were equally weighted. Reverse-complement augmentation was applied to all input sequences.

Following pretraining, we fine-tuned MethylAI on human data, generating four distinct models: one on the complete human dataset, one on ENCODE data, one on human atlas data, and one on HEK-293T cell line data. For the complete dataset, the model was initialized with pre-trained weights and trained for three epochs. Fine-tuning used the AdamW optimizer with a learning rate of 1×10^−4^ and a batch size of 300 (50 per GPU across 6 GPUs). A linear warm-up from 0 to 1×10^−4^ was applied over the first epoch, followed by a constant rate in the second epoch, and cosine decay from 1×10^−4^ to 1×10^−7^ in the third. Reverse-complement augmentation was applied to all sequences, and cell-type-specific hypomethylated regions were resampled to improve prediction accuracy.

For the ENCODE, human atlas, and HEK-293T-specific models, the following adjustments were made: a new output block was generated for each, with its output dimension tailored to the size of the corresponding dataset (i.e., S × 5). During fine-tuning, the learning rate was set to 1×10^−4^ for the input and multi-scale convolution blocks, and to 6×10^−4^ for the newly initialized output block. Additionally, for the human atlas dataset, a 5× oversampling strategy was applied to CpG sites located within cell-type-specific hypomethylated regions.

#### CpG embedding visualization

The output from the adaptive average pooling layer within the output block of MethylAI was extracted as a numerical vector representing the CpG embedding for each input DNA sequence. To visualize and analyze these embeddings, we performed dimensionality reduction using the multiscale PHATE algorithm^36^. The resulting low-dimensional representations were then used to examine and visualize the distribution of CpG sites originating from distinct genomic elements including CpG islands, CpG shore, CpG shelf, open sea, intergenic, exon, LINE, SINE and LTR.

### Evaluation and comparison with other models

#### Model evaluation

The validation set was used for model optimization and hyperparameter selection of MethylAI. The test set was employed for final evaluation and comparison with INTERACT and HyenaDNA. Prediction accuracy was assessed using the Pearson correlation coefficient (PCC) and Spearman correlation coefficient (SCC), calculated between predicted and observed methylation values across all CpG sites in the validation/test sets.

#### INTERACT model training

We obtained the INTERACT^12^ source code from the GitHub repository (https://github.com/LieberInstitute/INTERACT). The model output dimension was modified to match our completed human dataset (i.e. 7,870). The model was then trained from scratch on the human DNA methylation dataset for five epochs using the AdamW optimizer with a learning rate of 1.76×10^−4^ (as recommended in the GitHub repository) and a batch size of 300 (50 per GPU across 6 GPUs). A linear warm-up from 0 to 1.76×10^−4^ was applied during the first epoch, followed by a constant rate in the second epoch. For epochs 3-5, we used the *CosineAnnealingWarmRestarts* scheduler, decreasing the rate from 1.76×10^−4^ to 1.76×10−7 in epoch 3, and repeating this cycle over epochs 4-5. Reverse-complement augmentation was consistently applied during training.

#### HyenaDNA model fine-tuning

We obtained the pretrained HyenaDNA^13^ model (HyenaDNA-small-32k-seqlen) and its weights from Hugging Face (https://huggingface.co/LongSafari/hyenadna-small-32k-seqlen). To adapt it for DNA methylation prediction, we loaded the pre-trained model using the AutoModel class, extracted embeddings from the last token of the “last_hidden_state” output, and appended an output block identical to that used in MethylAI (linear layer → ELU activation → linear layer → sigmoid activation). The model was then fine-tuned on the human DNA methylation dataset for 3 epochs using AdamW with a learning rate of 1×10^−4^ for the HyenaDNA layers and 1×10^−3^ for the output block. Due to GPU memory constraints, the batch size was set to 48 (8 per GPU across 6 GPUs). We employed the same learning rate scheduler and reverse-complement augmentation strategy as used for MethylAI fine-tuning.

### Sequence attribution score and motif analysis

#### Nucleotide-resolution sequence attribution score

We computed nucleotide-resolution attribution scores using DeepSHAP, an implementation of the DeepLIFT algorithm^14^. Sequences centered on representative CpG sites (see below) were used as input to calculate base-resolution attribution scores, with the smoothed site methylation level of the corresponding sample serving as the prediction target.

#### Selection of representative CpG site

For lung tissue, to enrich for CpG sites associated with genes and CpG islands (CGIs), we randomly selected 10,000 CpG sites from each of the following genomic contexts: CpG shore, CpG shelf, open sea, core promoter (TSS ± 200 bp), extended promoter (TSS ± 1500 bp), 5’UTR, 3’UTR, exon, intron, and intergenic regions. Redundant sites were removed after merging. For CGIs, we selected the CpG closest to the center of each CGI shorter than 1 kb.

Top 1000 uniquely unmethylated regions (UURs) for each of 39 cell types were obtained from the human atlas dataset^24^. Genome-wide hypomethylated regions were defined as regions containing five or more consecutive CpGs with a smoothed methylation level < 0.25. For UURs and genome-wide hypomethylated regions, representative CpGs were selected as follows: for regions smaller than 1 kb, the CpG closest to the center was chosen; for larger regions, non-overlapping 1 kb windows were defined, and the CpG nearest the center of each window was selected.

#### Dinucleotide analysis

To analyze dinucleotide attribution scores, we first computed, for each input DNA sequence, the mean and total attribution scores of all 16 dinucleotide combinations within the ±1 kb window centered on the target CpG. These values were then averaged across all sequences to obtain the average mean and total attribution score per dinucleotide combination per input sequence.

#### Motif attribution and activation score

We obtained pre-computed predicted transcription factor motif sites for the human reference genome (hg38) from the UCSC database in BigBed format (https://hgdownload.soe.ucsc.edu/gbdb/hg38/jaspar/JASPAR2024.bb). This dataset includes genomic positions, motif identifiers, motif names, and a motif match score calculated as

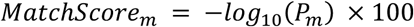

where the *Pm* indicates the significance of the position-specific motif profile match. We retained motifs with a motif match score > 400 (equivalent to p-value < 1×10^−4^) located within 1 kb of a representative CpG site.

The motif attribution score was defined as the mean absolute attribution score across all nucleotide positions within the motif match.

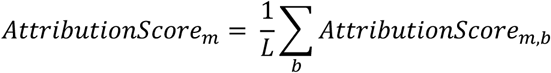

where *AttributionScore_m,b_* indicates the base pair attribution score for motif *m* and the base in position *b* in each motif site, *L* is the length of motif site.

The motif activation score (MAS) was calculated as the product of the motif match score and the motif attribution score.

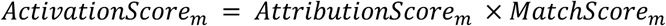

#### Normalized motif activation score

To enable cross-cell-type comparison of the MAS of TF motifs to UUR hypomethylation, we defined a normalized MAS (nMAS). For each TF, the total motif activation score (MAS) was summed across all UURs in a given cell type and normalized by the number of UURs in that cell type. The formula is as follows:

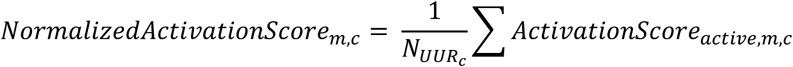

where *ActivationScore_active,m,c_* is motif activation score of an active motif for TF motif m and in cell-type *c*, *N_UURc_* is number of UURs in a given cell type, *NormalizedActivationScore_m,c_* is the normalized motif activation scores of a TF motif *m* in cell-type *c*.

#### Identification of active motifs

For CGIs, motifs with an absolute attribution score > 0.02 were classified as active. For UURs and genome-wide hypomethylated regions, motifs with attribution scores < −0.02 were defined as active, reflecting their potential role in maintaining hypomethylation.

For UURs and genome-wide hypomethylated regions, we applied two additional processing steps to reduce false positives. First, to minimize the influence of methylation prediction inaccuracies, we excluded representative CpG sites where the absolute difference between MethylAI-predicted and observed methylation values exceeded 0.2 (affecting ~2.5% of sites on average). Second, when a motif fell within ±1 kb of two representative CpGs (occurring when the represent CpG sites were < 2 kb apart), we retained the minimum attribution score to avoid double-counting and to reflect the strongest hypomethylation-contributing signal, ensuring each motif site was assigned a unique attribution and activation score.

### Motif analysis and experimental validation in HEK-293T cells

#### Construction of CTCF and JUN family knockout cell lines

The sgRNA oligonucleotides targeting CTCF was designed according to the Zhang Lab protocol. Oligos were annealed and cloned into the BsmBI-linearized LentiCRISPR-v2 vector (Addgene #52961). Construct was verified by Sanger sequencing.

HEK-293T cells were transfected with the resulting plasmids using Lipofectamine 3000 (Invitrogen L3000015). At 24 hours post-transfection, cells were selected with 1 μg/mL puromycin for 5-7 days to obtain polyclonal populations: HEK-293T-Nontargeting^CK^, HEK-293T-CTCF^KO^.

The following sgRNA target sequences were used:

5’-AAAAAGCTTCCGCCTGATGG-3’ (sgNontargeting)

5’-GAGCAAACTGCGTTATACAG-3’ (sgCTCF)

Knockout efficiency was validated by genomic DNA extraction (QIAGEN #69504), PCR amplification of regions flanking each sgRNA target, and Sanger sequencing to assess indel patterns. Total RNA was also extracted (YEASEN #19221ES50) to quantify mRNA expression of the targeted genes.

#### WGBS library preparation and sequencing

Cells (HEK-293T, HEK-293T-Nontargeting^CK^ and HEK-293T-CTCF^KO^) were harvested by centrifugation, and genomic DNA was extracted using a standard DNA extraction kit. One microgram of DNA was sonicated into fragments of 200-500 bp. Fragments were end-repaired, A-tailed, and ligated to barcoded adapters. Bisulfite conversion was performed using the Hieff Superfast DNA Methylation Bisulfite Kit (YEASEN 12225ES50), during which unmethylated cytosines were deaminated to uracils. Desulfonation was carried out to remove sulfonation groups, and converted DNA was amplified using uracil-tolerant polymerase (NEB M0597S). Libraries were quantified with Qubit, and size distribution was assessed using the Qseq system. Paired-end sequencing was performed on the Illumina platform at 30× coverage to ensure sufficient depth for methylation calling.

#### WGBS processing and MethylAI fine-tuning

WGBS data were processed using Bismark^37^. Raw FASTQ files were first trimmed with Cutadapt to remove adapter sequences and low-quality bases. Resulting clean reads were quality-checked with FastQC. The human reference genome (hg38) was indexed using bismark_genome_preparation, followed by alignment with bismark to generate BAM files. Duplicate reads were removed using deduplicate_bismark, and raw site methylation levels at all CpG sites were extracted using bismark_methylation_extractor. After, smoothed site levels and regional methylation levels were obtained as previously described. A HEK-293T-specific MethylAI model was generated by fine-tuning the pretrained MethylAI for three epochs using the parameters described in MethylAI pre-training and fine-tuning Section.

#### Active motif identification in genome-wide hypomethylated regions

Genome-wide hypomethylated regions were defined as regions containing five or more consecutive CpG sites with smoothed methylation levels < 0.25. Representative CpG sites were selected for each region using the criteria described in Section 3.1. Motif attribution scores were calculated for all motif instances within ±1 kb of these CpG sites. Given the hypomethylated status of these regions, motifs with attribution scores < −0.02 were classified as active motifs.

#### Interactions of transcription factor with epigenetic factors

Protein-protein interaction (PPI) data were obtained from the STRING database (https://cn.string-db.org/), retaining edges with a confidence score ≥ 0.4. Active motifs of TFs were ranked by the number of hypomethylated regions in which they occurred, and the top 30% were designated as top TFs. Enrichment of interactions between these top TFs and epigenetic factors was assessed by calculating odds ratios, using all TF of active motifs identified in hypomethylated regions as the background set.

#### TF occupancy and histone modification profiling

Publicly available ChIP-seq data for TFs and histone modifications in HEK-293/HEK-293T were obtained from ENCODE. To assess TF occupancy, active and inactive motif instances were intersected with TF ChIP-seq peaks; motifs overlapping a peak were considered TF occupied. For histone modification analysis, genomic intervals surrounding active or inactive motifs for CTCF, JUN, and GATA6 were merged using BEDTools separately. Signal matrices for active/inactive sets were generated with deepTools computeMatrix over ±1 kb windows, and profiles were visualized with plotProfile.

### In Silico Variant and Motif Knockout Prediction Using MethylAI

#### Enhancing Genotype-Tissue Expression (eGTEx) mQTL dataset

We obtained summary statistics for both regular and conditional mQTLs from the GTEx Portal^29^ (https://gtexportal.org/home/datasets) across seven tissues: breast mammary tissue, colon transverse, lung, skeletal muscle, ovary, prostate, and testis. These tissues were selected due to their strong concordance with samples in our ENCODE dataset. We retained significant mQTL-CpG pairs (nominal P < 1×10^−5^) where the distance between the mQTL and the target CpG (mCpG) was within 9,216 bp—half the input sequence length of MethylAI—to ensure predictable context.

#### GWAS Catalog variant datasets

Files of GWAS Catalog associations data and all traits to EFO mappings were downloaded from https://www.ebi.ac.uk/gwas/docs/file-downloads in August 2025. To enhance the focus of the analysis and reduce the multiple testing burden a priori, we prioritized traits belonging to the following EFO parent categories: Metabolic-related (Lipid or lipoprotein measurement, Metabolic disorder),Immune-related (Inflammatory measurement, Immune system disorder), Hematological-related (Hematological measurement, Neurological disorder, Anthropometric-related (Body measurement), Cancer, Cardiovascular-related (Cardiovascular measurement, Cardiovascular disease), Digestive-related (Digestive system disorder), and Biological process. Subsequently, we identified 5,600 traits linked to top associations located within active motifs of 52 selected tissues for downstream analysis.

#### Prediction of variant effects

For in silico mutagenesis analysis of mQTLs and GWAS variants, we used MethylAI to predict smoothed site methylation levels for both reference and alternative alleles. The predicted effect of each variant was calculated as the difference between alternative and reference methylation levels (Δmethylation = alt − ref). To assess directional accuracy, we compared the sign of Δmethylation with the signed statistical slope of the original mQTL association.

#### In Silico motif knockout

To simulate motif knockout, we performed in silico deletion by removing nucleotides corresponding to the motif (i.e., replacing each specific motif sequence with a null string). The input sequence was then recentered to maintain the target CpG at the center of the 18,432 bp window. MethylAI was used to predict smoothed methylation levels after motif deletion, and the effect was quantified as the difference between knockout and wild-type sequences (Δmethylation = KO − Control).

#### Predicted Directional Effect of mQTL Variants

We collected the mQTLs, including conditionally independent mQTLs, within a ±500 Kb window of each CpG locus from the Enhancing GTEx (eGTEx) project [PMID: 36510025], integrating data across seven eGTEx tissues (breast mammary tissue, colon transverse, lung, skeletal muscle, ovary, prostate, and testis), given that these tissue types show higher concordance with those represented in ENCODE-based fine-tuned models. Our analysis focused on CpG sites located in cis that exhibited a significant association with an mQTL (FDR < 0.05), which termed mCpGs. Then we filtered for significant mQTLs (defined at a nominal P< 1×10^−5^), centered the input sequence on each mQTL, and recorded the methylation level of all associated mCpGs for both the reference and alternative nucleotides of the causal variant. For each mQTL-mCpG pair, we computed variant effect sizes, defined as the difference between alternative and reference predictions. To validate the predictions, we evaluated the agreement between the direction of the model-predicted variant effect and the signed statistical slope of the mQTL association. The accuracy of our model’s directional predictions for mQTLs was evaluated specifically on its identified active motifs. This was compared to the baseline accuracy calculated for mQTLs located within (i) the ±1 kb flanking regions of the mCpGs (including inactive motifs, non-motif, and random regions), and (ii) arbitrary regions across the genome without distance constraints. For the random regions within the ±1 kb flanking regions and thearbitrary regions (without distance constraints), mQTLs were randomly sampled to match the number in active motifs.

#### Evolutionary conservation analysis

To compare the evolutionary conservation around mCpG sites, we calculated the mean 470-way mammalian phyloP scores within merged active motifs, inactive motifs, and non-motif regions located in the ±1 kb flanking regions. Regions with phyloP scores less than −1 were defined as fasting-evolution, those with scores between −1 and 1 as neutral evolution, and those with scores greater than 1 as evolutionarily conserved.

### Enrichment of GWAS variants in tissue-specific active motifs

We extracted trait-associated variants that were located within either active or inactive motifs, in any tissue. To determine the significance of enrichment, Fisher’s exact test was used to evaluate if top associations across various trait categories were significantly enriched in tissue-specific active motif regions. A false discovery rate (FDR) of < 0.1 was considered statistically significant. Due to similarities in the PWM matrices of certain motifs, a single binding site may match multiple motifs. In such cases, the active motif with the highest match score is selected as the representative for presentation. The match score between the DNA sequence containing the SNP’s risk allele and this assigned motif was then recalculated using the PWMScan.

## Code availability

The code for the MethylAI model can be accessed at https://github.com/Yu-Lab-Genomics/MethylAI.

## Author contributions

F.Y. and F.C conceived the project and designed the algorithm. F.C implemented the algorithm. Q.Z. constructed the multi-species DNA methylation dataset. Shiwei Z. performed mQTL and GWAS variants analysis. J.L. performed WGBS and CRISPR experiment of HEK-293T cell line. F.C., Q.Z., Shiwei Z. and Z.Z, performed computational experiments and contributed to data interpretation. Siyun Z. developed the companion website. F.Y., F.C., Q.Z., Shiwei Z. and J.S. wrote the manuscript with input from all authors. F.Y., Y.Z., X.F. and Y.G. supervised and directed the study.

## Acknowledgements

We are grateful to members of the Yu laboratory and numerous colleagues for valuable comments and suggestions. This work was supported in part by the grant 2023YFF1204701 from the National Key R&D Program of China, grant 2024B1515020080 from Guangdong Basic and Applied Basic Research Foundation, grants 32470634 and 32460155 from the National Natural Science Foundation of China, grant KY012023362 from the Talent Research Funding Project of Guangdong Provincial People’s Hospital, grants GZNL2024A01003 and GZNL2023A02002 from the Major Project of Guangzhou National Laboratory, grants 202302AD080004, 202403AC100002 from Key R&D Program of Yunnan and grant 2024YNLCYXZX0195 from Clinical Medical Center for Cardiovascular Disease of Yunnan Province. We acknowledge the Data Science Platform of Guangzhou National Laboratory and the Bio-medical Big Data Operating System (Bio-OS) for technical support and for providing access to the computational resources essential to this study.

